# Waste oil substrates reshape the black soldier fly larval gut microbiome and biomass composition during bioconversion

**DOI:** 10.64898/2026.06.09.731207

**Authors:** Reese Saho, Duy Trinh, Erin Kojima, Tong Wang, Charity Owings, Zachary Burcham

## Abstract

Black soldier fly larvae (BSFL) are generalist decomposers with promise for converting agricultural and food-processing by-products into value-added bioproducts, but BSFL performance on lipid-rich waste oil streams and the role of the gut microbiome in this process remains unclear. Here, we evaluated BSFL bioconversion of a standard chicken feed diet supplemented with three chemically distinct waste oils: acidulated vegetable oil (AVO), pork grease (PG), and used cooking oil (UCO). Larval performance, bioconversion rate, gut microbiome composition, total protein and fat content, and fatty-acid profiles were measured across bioconversion. Larval age was a major driver of gut microbiome structure, but waste oil supplementation further reshaped community membership and structure, with the strongest diet-associated effects occurring during early-to-intermediate bioconversion. Most differentially abundant taxa were members of the baseline core gut community, suggesting that oil supplementation primarily altered dominance patterns among resident taxa. PG and UCO supported larval growth and bioconversion performance comparable to the chicken feed control, whereas AVO reduced bioconversion rate and showed weaker growth outcomes. Oil supplementation also increased larval fat content, reduced protein content, and shifted fatty-acid profiles toward the corresponding oil feedstocks, although larval biomass composition remained shaped by basal diet and host or microbial metabolism. These findings show that selected lipid-rich waste streams can support efficient BSFL bioconversion while restructuring resident gut microbiome members that may tolerate, metabolize, or indirectly respond to oil-associated conditions, contributing to substrate-dependent changes in larval lipid accumulation and fatty-acid composition.

**IMPORTANCE:** Agricultural and food-processing systems generate large amounts of lipid-rich by-products that are difficult to manage using conventional waste-valorization approaches. Black soldier fly larvae (BSFL) offer a biological route for recovering nutrients from these materials, but efficient conversion depends on interactions among substrate chemistry, larval physiology, and the gut microbiome. This study shows that selected waste oil streams can support larval growth while restructuring resident gut microbial communities and altering larval fatty-acid composition. These findings are important for agricultural biotechnology because they frame BSFL production as a host-microbiome bioconversion system rather than simply an insect-based waste-reduction process. Understanding how gut microbes respond to chemically distinct lipid wastes can guide substrate selection, pretreatment, and microbiome-informed optimization strategies for converting underutilized agricultural and food-processing residues into value-added bioproducts for circular agricultural systems.

## INTRODUCTION

The black soldier fly larva (BSFL), *Hermetia illucens* (Linnaeus) (Diptera: Stratiomyidae), is a generalist decomposer with high bioconversion efficiency and growing importance in circular bioeconomy systems (1, 2). BSFL can develop on diverse organic waste streams (3–6), including agricultural residues, food-processing by-products, and post-consumer food wastes, converting low-value materials into insect biomass that can be used as alternative feedstuff for livestock and frass with biofertilizer potential. BSFL are also being investigated for their potential role in transforming more recalcitrant materials such as lignin- and plastic-associated substrates (7–10). Because BSFL bioconversion can recover nutrients from wastes while reducing reliance on conventional disposal routes, BSFL production is increasingly investigated as a sustainable approach to organic waste valorization. The ability of BSFL to use diverse resources is linked, in part, to the gut microbiome, which contributes to nutrient breakdown and acquisition (11). For example, conventionally reared larvae attain greater biomass than axenic larvae, indicating that gut microbes can contribute to larval growth and development (12, 13). The BSFL gut microbiome is also highly responsive to diet and shifts in composition when larvae are reared on different substrates, including anthropogenic waste streams (14–17). These diet-associated shifts suggest that substrate chemistry can impose selective pressures that favor gut community members able to tolerate, metabolize, or indirectly benefit from specific substrate components. Despite this plasticity, BSFL (18, 19), and Stratiomyidae more broadly (20), may retain a core set of gut-associated microbes, including *Dysgonomonas*, *Morganella*, *Ignatzschineria*, *Providencia*, and *Enterococcus*. These taxa may contribute to host fitness through nutrient acquisition, reproductive success, or competitive exclusion of pathogens (21), although their specific ecological roles remain understudied in BSFL. Therefore, understanding how resident and diet-responsive gut microbes shift in response to chemically complex waste substrates is important for improving BSFL-based bioconversion systems.

Lipid-rich waste streams represent an important but chemically variable class of organic by-products. The production and use of culinary and food-associated oils generate low-value residues with high caloric content, including acidulated oils from vegetable-oil refining and used cooking oils from repeated heating. For example, refining crude vegetable oil produces soapstock, which can be converted to acidulated oils containing high proportions of long-chain free fatty acids (FFAs) (22). During use, heating plant- and animal-based cooking oils promotes lipid hydrolysis, oxidation, and polymerization, generating FFAs, polar compounds, aldehydes, ketones, and polymerized acylglycerols (23, 24). These chemical changes reduce oil quality, impact digestibility and nutritional value, and make waste oils more challenging to process than unmodified oil substrates (25). Conventional management of waste oils, such as incineration, landfilling of residual sludge, and chemical treatment, often limits valorization potential and contributes to environmental contamination in wastewater systems, soils, and waterways (26). To recover value from these lipid-rich residues, waste oils are commonly converted into biofuels, particularly biodiesel. However, this route can be constrained by feedstock variability, high FFA content, pretreatment requirements, and competition for available waste-oil supply (23, 27). These limitations have increased interest in alternative biological strategies for recovering value from lipid-rich wastes. A previous study directly tested BSFL conversion of canteen-derived oil-separator waste, but the pure oil-waste diet severely limited larval performance (14). Larvae fed this substrate showed negligible substrate reduction, no pupation, inhibited biomass gain by 85%, and up to 96% mortality, likely attributed the increasingly viscous fats, oils, and grease impaired larval movement, feeding, and potentially respiration through physical blockage of spiracles. However, this study also showed that oil-rich waste can alter the BSFL gut microbiome, with oil-and food-waste diets diverging from chicken feed and enriching taxa such as *Morganella* and *Acinetobacter* during development. These findings suggest that oil-rich wastes can influence both BSFL performance and gut community structure, but the poor performance on a pure oil-separator waste diet limits conclusions about whether chemically diverse waste oils can be used effectively as supplements within nutritionally balanced substrates.

In this study, we reared BSFL on a standard chicken feed diet with or without supplementation of three chemically distinct waste oils: acidulated vegetable oil, pork grease, and used cooking oil. These substrates represent different lipid waste chemistries, including free fatty acid-rich acidulated oil, more saturated animal-derived grease, and thermally altered cooking oil. We hypothesized that waste oil supplementation would reshape the BSFL gut microbiome in a substrate-specific manner, favor putative lipid-tolerant or lipid-associated taxa, and alter larval biomass composition by increasing lipid accumulation and shifting fatty-acid profiles toward those of the supplemented oils. To test this, we measured larval growth and bioconversion rate to assess performance on each oil-supplemented diet. We profiled larval gut microbiomes across bioconversion to identify temporal changes in community membership and structure. Finally, we measured endpoint larval protein, fat, and fatty-acid composition to determine how waste oil supplementation influenced the nutritional profile of harvested biomass.

## METHODS

### BSFL nursery phase

Twelve billets of BSF neonates (∼10,000 larvae per billet) were obtained from Fluker’s (Baton Rouge, Louisiana, USA). Each billet was used to inoculate 1 kg chicken feed (Purina^®^ Layena^®^; St. Louis, Missouri, USA) adjusted to 60% moisture within a 5.67 L plastic shoe box (Sterilite^®^) with an aerated lid. Each replicate was reared for four days in an I-36VL environmental growth chamber (Percival Scientific™; Perry, Iowa, USA) set at 30°C, 70% RH and a 12:12 light:dark cycle. At the end of the nursery phase, BSFL from each treatment were sieved through a 6.35 mm gold mining pan and weighed.

### BSFL bioconversion phase

Three waste oils were tested in the bioconversion phase: acidulated vegetable oil (AVO), pork grease (PG), and used cooking oil (UCO). The waste oil substrates were obtained/produced from Feed Energy Company (Pleasant Hill, IA, USA). AVO was produced from soapstock generated during vegetable oil refining. PG was a non-food-grade animal fat produced from pork trimmings collected from meat-packing facilities. UCO consisted of restaurant-derived used cooking oil. Each treatment was prepared as a 10% oil inclusion by weight at 60% moisture using the following formula: 300 g chicken feed (CF), 100 g oil, 600 g distilled water. The negative control diets were made with 400 g CF and 600 g distilled water. Each treatment and control were replicated in triplicate, and all were contained within 5.67 L plastic shoe boxes (Sterilite^®^). Sieved larvae from each nursery replicate were randomly assigned to a treatment and one of three replicate bioconversion bins per treatment for the bioconversion phase. Larvae were maintained in the bioconversion phase for seven days under the same environmental conditions as the nursery phase. Three random larvae from each replicate bioconversion bin were obtained on the last nursery day after sieving (i.e., bioconversion day 0), as well as on bioconversion days two and seven for microbiome analysis. At the end of the seven-day bioconversion phase, larvae from each replicate were parboiled at ∼100°C for 1 min and frozen at −20°C for downstream fatty acid composition analysis and total fat and protein quantification.

### Statistical analysis of BSFL growth and bioconversion traits

The following trait data for BSFL were obtained: total larval mass, residue mass, net larval gain, diet mass loss, and bioconversion rate for both the nursery phase and bioconversion phase. Bioconversion rate (BCR) was calculated using the following formula: (ending larval mass/initial dry diet mass) × 100. Overall net larval gain (e.g., from neonate to harvest) was also obtained. Data were analyzed using one-way ANOVA with a post-hoc Tukey’s HSD pairwise comparisons test.

### BSFL gut dissection, DNA extraction, 16S rRNA gene library preparation, & amplicon sequencing

On each bioconversion day (0, 2, and 7), 3 larvae per bioconversion bin per diet were collected for gut microbiome analysis, resulting in a total of 9 larval replicates per diet per day and 27 larval replicates per diet. Larvae were surface sterilized using 70% ethanol before whole guts were dissected. Guts were frozen at −80°C until all samples were collected. Prior to DNA extraction, 250uL of 1X autoclave sterilized phosphate-buffered saline was added to thawed guts. The mixture was homogenized with a sterile pestle until a homogenous solution formed. Six blank negative extraction control samples and a positive ZymoBIOMICS Microbial Community Standard (Cat. #: D6300) control sample were included during DNA extraction. Extraction was conducted using the KingFisher Flex: MagMAX Microbiome Soil extraction protocol. DNA was quantified using a Quant-iT PicoGreen dsDNA assay (Cat. #: P7589). DNA was then prepared for 16S rRNA gene library sequencing following the Earth Microbiome Project (EMP) 16S V4 rRNA gene amplification protocol with the 515F-Parada (FWD: 5’GTGYCAGCMGCCGCGGTAA) and 806R-Apprill (REV: 5’GGACTACNVGGGTWTCTAAT) primers. Libraries were pooled to an equimolar ratio and sequenced on an Illumina NEXTSeq to generate 2 x 300bp paired-end sequences at the UTK Genomics Core.

### Microbiome data pre-processing

Sample processing and filtration was conducted using the QIIME2 Amplicon program suite (v. 2026.1) (28). All programs were used with default settings unless otherwise described. Sequence reads were demultiplexed, trimmed to 240 bp to remove low quality tails, and denoised using the paired-end DADA2 plugin (v. 1.30.0) (29). This resulted in 37,918,915 total reads and 336,291.5 median reads per sample. To identify putative contamination, the QIIME2 decontam-identify plugin (v. 1.22.0) (30) was utilized with the prevalence-based method. Reads with a decontam score of < 0.2 were removed from analysis. Approximately 327,772 (0.86%) of reads were identified as putative contaminants, leading to a total of 37,591,143 reads remaining. A phylogenetic tree was generated using the QIIME2 fragment-insertion plugin (28) into the SILVA 128 99% reference tree (31). Taxonomic information was assigned to reads using the QIIME2 classify-sklearn plugin (32) to train a naïve Bayes classifier on the GreenGenes2 99% reference database (33) as a reference. Amplicon sequence variants (ASVs) were then filtered to remove any reads that correspond to mitochondria or chloroplasts using the QIIME2 filter-table plugin (28). Taxonomic assignments were used to generate relative abundance charts in R’s (v 4.5.1) ggplot2 package (v. 4.0.0).

### Microbiome community analyses

Alpha rarefaction curves were generated using the QIIME2 alpha-rarefaction plugin (28) to evaluate the appropriate depth to normalize the number of reads in each sample for diversity metric calculations. To analyze differences in microbial communities, alpha- and beta-diversity metrics were generated using the QIIME2 core-metrics-phylogenetic plugin (28). Alpha- (Faith’s phylogenetic diversity, Pielou’s evenness, Shannon’s diversity, and ASV richness) and beta-diversity (weighted and unweighted UniFrac) metrics were calculated at a rarefaction depth of 34,000 reads to maximize sample retention and ensure adequate representation of the microbial communities within each sample. All negative controls fell below this threshold, leading to their removal from future analysis. To statistically compare microbial community diversity and calculate the proportion of variance due to specific covariates, the QIIME2 adonis plugin was used to conduct a multivariate PERMANOVA (perm = 999) (34) on beta diversity distance matrices. The QIIME2 beta-group-significance plugin was used to conduct pairwise PERMANOVA analyses across bioconversion day and diet (perm = 999). After stratifying by bioconversion day, bioconversion bin number was included as a co-variate to account for potential batch effects. Statistical analysis of the alpha-diversity metrics was calculated in R (v. 4.5.1) (35) using a Kruskal-Wallis test (v. 4.5.1) followed by a post-hoc Dunn’s test (v. 1.3.6) to determine significant pairwise differences. To determine differentially abundant taxa relative to the control chicken feed diet with no oil addition, the QIIME2 ANCOMBC2 plugin (v. 2.4.0) was conducted on non-rarefied samples using the structural zeros method. Differential abundance heatmaps were generated using ggplot2 in R. A core features analysis was performed using the QIIME2 feature-table plugin (28). Taxa were filtered into 50%+, 80%+, or 100% prevalence. Any taxa that were 80%+ prevalence and detected in each diet group were considered core taxa.

### BSFL total fat and protein quantification

The Folch extraction was utilized to extract lipids (36). Approximately 10 g of ground frozen BSFL biomass from each treatment was mixed with 50 mL of chloroform:methanol (2:1 v/v) and stirred for 30 minutes at room temperature before filtration with Whatman Grade 1 filter paper (Maidstone, UK) to recover liquid fraction. The solid fraction was then extracted for a second and a third time. The liquid fractions were combined and roto-evaporated at 60°C and 337 mbar (80 RPM) for 2 hours to recover total lipid. The mass of the final lipid pool was used to calculate total fat content. Defatted solid fractions were dried in the oven at 50°C overnight and saved for further analyses. Approximately 50 mg of defatted sample was used for total nitrogen content by Dumas method (37) using the vario MAX C/N Analyzer (Elementar Americas, Ronkonkoma, NY 11779). A conversion factor of 6.25 (38) was applied to determine total protein content of the sample. Data were analyzed using one-way ANOVA with a post-hoc Tukey’s HSD pairwise comparisons test.

### BSFL fatty acid composition analysis

Recovered lipids from BSFL after defatting and raw waste oil substrates were converted to fatty acid methyl esters (FAMEs) using acid catalysis (39). Briefly, approximately 20 µL of lipids were mixed with 2 mL of 4% H_2_SO_4_ in methanol (v/v) in a glass tube with a Teflon-lined cap and allowed for reaction to occur overnight in an oven at 80°C. After reaction, 2 mL of water and 2 mL of hexanes were added to allow for separation. The hexane fraction was collected, water washed and saved for gas chromatography analysis. Fatty acid composition was determined using capillary gas chromatography GC-2010 (Shimadzu, Kyoto, Japan) with a flame ionization detector and capillary HP-88 high polarity column (100 m × 0.25 mm × 0.20 µm) (Agilent Technologies, Santa Clara, CA, USA) as described by Angeletti et al. (40). The oven temperature program was as follows: initial temperature of 100°C and hold for 2 min, then ramp up to 230°C at 10°C/min and hold for 5 min. Total analysis time was 20 min. The injector and detector temperatures were 180°C and 250°C, respectively. The split ratio was 10:1, the sample volume was 1 μL and carrier gas was helium. Total flow was 15.6 mL/min and purge flow was 3.0 mL/min with a pressure of 110.9 kPa. Signal peaks were identified by referencing the retention times in standard 15A (Nu-Check Prep, Elysian, MN, USA) for C16:0 to C20:1 and commercial coconut oil for C8:0 to C14:0. Fatty acid composition of a sample was normalized calculated by dividing the identified peak area by the total peak area. A biplot fitting fatty acid relative abundances to their contribution to the lipid profile was calculated using the vegan R package (v. 2.7.1) (41).

### Data availability

Raw FASTQ files are publicly available on the NCBI Sequence Read Archive (SRA) under BioProject PRJNA1471181. Analysis scripts and metadata are provided in the Burcham Lab GitHub project repository (https://github.com/BurchamLab/2026_bsfl_waste_oil).

## RESULTS

### Larval Age is a Primary Driver of Gut Microbiome Differentiation

Larval age explained the largest proportion of variation in the presence and absence of gut microbial taxa (i.e., community membership) (unweighted UniFrac R^2^ = 0.25; p = 0.001), followed by diet (unweighted UniFrac R^2^ = 0.17; p = 0.001; **Figure 1A-B**). A similar trend was observed when accounting for the taxon relative abundances within communities (i.e., community structure), but with less variation assigned to both age (weighted UniFrac R^2^ = 0.16; p = 0.001) and diet (weighted UniFrac R^2^ = 0.12; p = 0.001; **Figure S1**). Gut microbial communities diverged progressively over bioconversion, following a directional trajectory from day 0 through day 2 to day 7, with the greatest separation between the earliest and latest timepoints (unweighted UniFrac F = 14.86, q = 0.001; weighted UniFrac F = 12.89, q = 0.001; **Table S1**). Within-community alpha diversity metrics reinforced this temporal pattern (**Figure 1C-E**, **Table S2**). Phylogenetic diversity decreased significantly in the beginning stages of bioconversion (day 0 vs. 2), did not differ between days 2 and 7, and maintained the reduction at the end of bioconversion (day 0 vs. day 7). Evenness trended downward from day 0 to 2, significantly increased from day 2 to 7, but was not significantly different at day 7 compared to day 0. Shannon’s entropy showed a transient reduction at day 2 relative to both day 0 and day 7.

**Figure 1.**
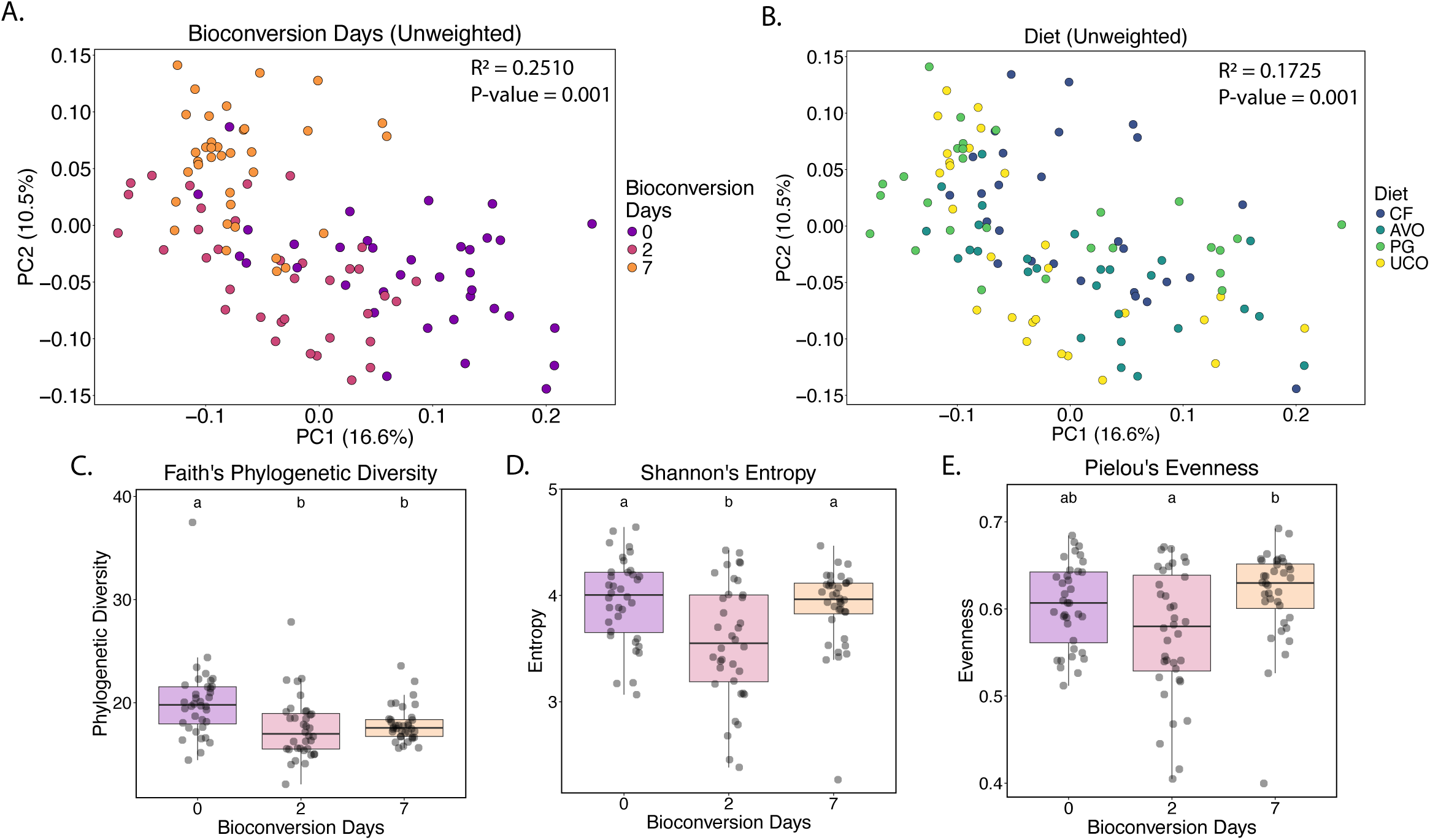
Larval age significantly impacts gut microbiome composition. (A-B) PCoA ordinations of unweighted UniFrac distances colored by (A) bioconversion day or (B) diet. CF = chicken feed control, AVO = acidulated vegetable oil, PG = pork grease, and UCO = used cooking oil. Data were analyzed using a multivariate PERMANOVA. (C-E) Boxplots demonstrating change in (C) Faith’s phylogenetic diversity, (D) Pielou’s evenness, and (E) Shannon’s entropy across bioconversion days. Data were analyzed using Kruskal-Wallis test with post-hoc Dunn’s tests. Letters indicate significant pairwise differences between days (q < 0.05).

### Waste Oil Substrates Progressively Reshape Gut Microbial Communities

Globally, gut microbial community membership (unweighted UniFrac R^2^ = 0.17; p = 0.001; **Figure 1B**) and community structure (weighted UniFrac R^2^ = 0.12; p = 0.001; **Figure S1**) differed among diets, and most diets supported distinct gut microbial communities (**Table S3**). The only exception was between UCO- and PG-fed larvae, which had similar community structures (**Table S3**). Because community composition also strongly varied across bioconversion days, subsequent analyses were stratified by day to investigate the temporal changes in diet-associated community composition throughout bioconversion. Although larvae were reared under identical conditions prior to oil supplementation, significant differences in community membership were observed between assigned diet groups at day 0 (unweighted UniFrac R^2^ = 0.14; p = 0.001; **Figure 2A**), but this did not extend to differences in community structure (weighted UniFrac R^2^ = 0.08; p = 0.12; **Figure S3**). Bin-level batch effects were also detected at day 0 and accounted for a larger proportion of variation in both community membership (unweighted UniFrac R^2^ = 0.32; p = 0.001) (**Figure 2A**) and community structure (weighted UniFrac R^2^ = 0.54; p = 0.001) (**Figure S3**). Despite this initial variation, diet-associated differences became more pronounced as bioconversion progressed.

**Figure 2.**
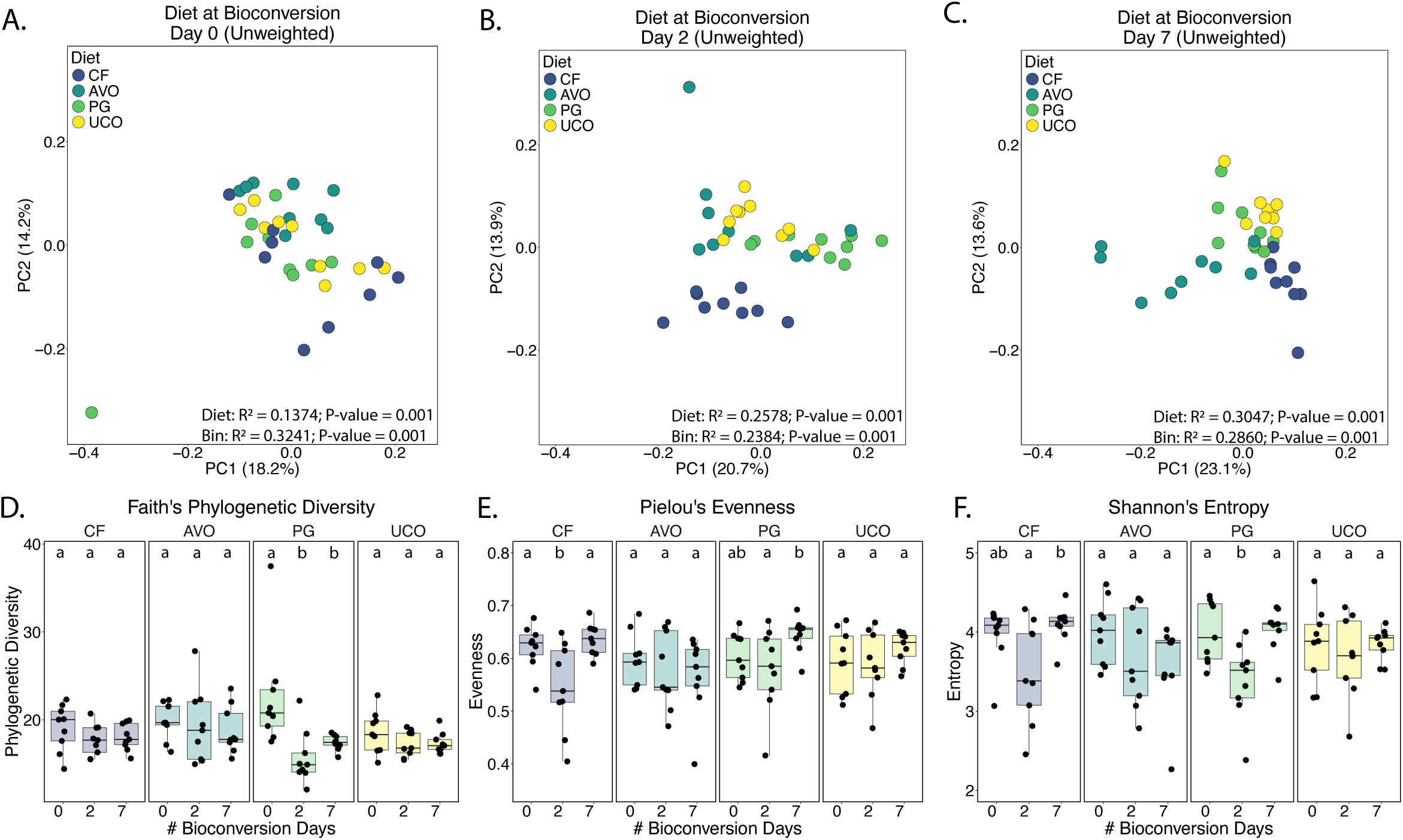
Diet leads to divergence of microbial communities over time. (A-C) PCoA ordinations of unweighted UniFrac distances colored by diet after 0 (A), 2 (B), or 7 (C) days of bioconversion. CF = chicken feed control, AVO = acidulated vegetable oil, PG = pork grease, and UCO = used cooking oil. Data were analyzed using a multivariate PERMANOVA. (D-F) Boxplots demonstrating change in (D) Faith’s phylogenetic diversity, (E) Pielou’s evenness, and (F) Shannon’s entropy across diet and bioconversion days. Data were analyzed using Kruskal-Wallis test with post-hoc Dunn’s tests. Letters indicate significance differences between days within each diet (q < 0.05).

Diet-associated separation became stronger over time for community membership, with diet explaining progressively more variation from day 0 to day 7 (unweighted UniFrac day 0 R² = 0.14; day 2 R² = 0.26; day 7 R² = 0.30; **Figure 2A-C**; **Table S4**). Bin-level effects were also detected across days and explained a similar proportion of community membership variation at each timepoint (unweighted UniFrac day 0 R² = 0.32; day 2 R² = 0.24; day 7 R² = 0.29; **Figure 2A-C**; **Table S4**). For community structure, diet explained the most variation at day 2 and remained a strong signal at day 7 (weighted UniFrac day 0 R² = 0.08; day 2 R² = 0.35; day 7 R² = 0.25; **Figure S3**; **Table S4**). Similar to with community membership, bin effects on community structure variation were strongest at day 0 but remained detectable at later timepoints (weighted UniFrac day 0 R² = 0.54; day 2 R² = 0.26; day 7 R² = 0.32; **Figure S3**; **Table S4**). Community phylogenetic diversity, evenness, and Shannon’s diversity were less consistently impacted by diet than community membership and community structure (**Figure 2D-F**; **Table S5**). Within individual diet treatments, PG-fed larvae had lower phylogenetic diversity on days 2 and 7 than on day 0 (**Figure 2D**). Evenness decreased in CF-fed larvae on day 2 relative to days 0 and 7, while PG-fed larvae showed higher evenness on day 7 (**Figure 2E**). Shannon’s diversity also decreased on day 2 in CF- and PG-fed larvae relative to days 0 and 7 (**Figure 2F**). Class-level relative abundance profiles showed that BSFL gut communities were consistently dominated by Actinomycetes, Bacilli, and Gammaproteobacteria across diets and time points (**Figure 3, Table S6**). However, the relative abundances of these classes shifted over time and differed among diets, particularly, at day 2 for the oil-supplemented diets and day 7 for the CF diet. These day 2 class-level shifts coincided with stronger diet-associated differences in both community membership and community structure. In CF-fed larvae, Actinomycetes proportionally increased at day 2, coinciding with reduced evenness and Shannon diversity. This proportional increase in Actinomycetes was not observed in the oil-supplemented diets, which instead showed proportional increases in Gammaproteobacteria and Bacteroidia. By day 7, all diet groups showed an increased proportion of Clostridia.

**Figure 3.**
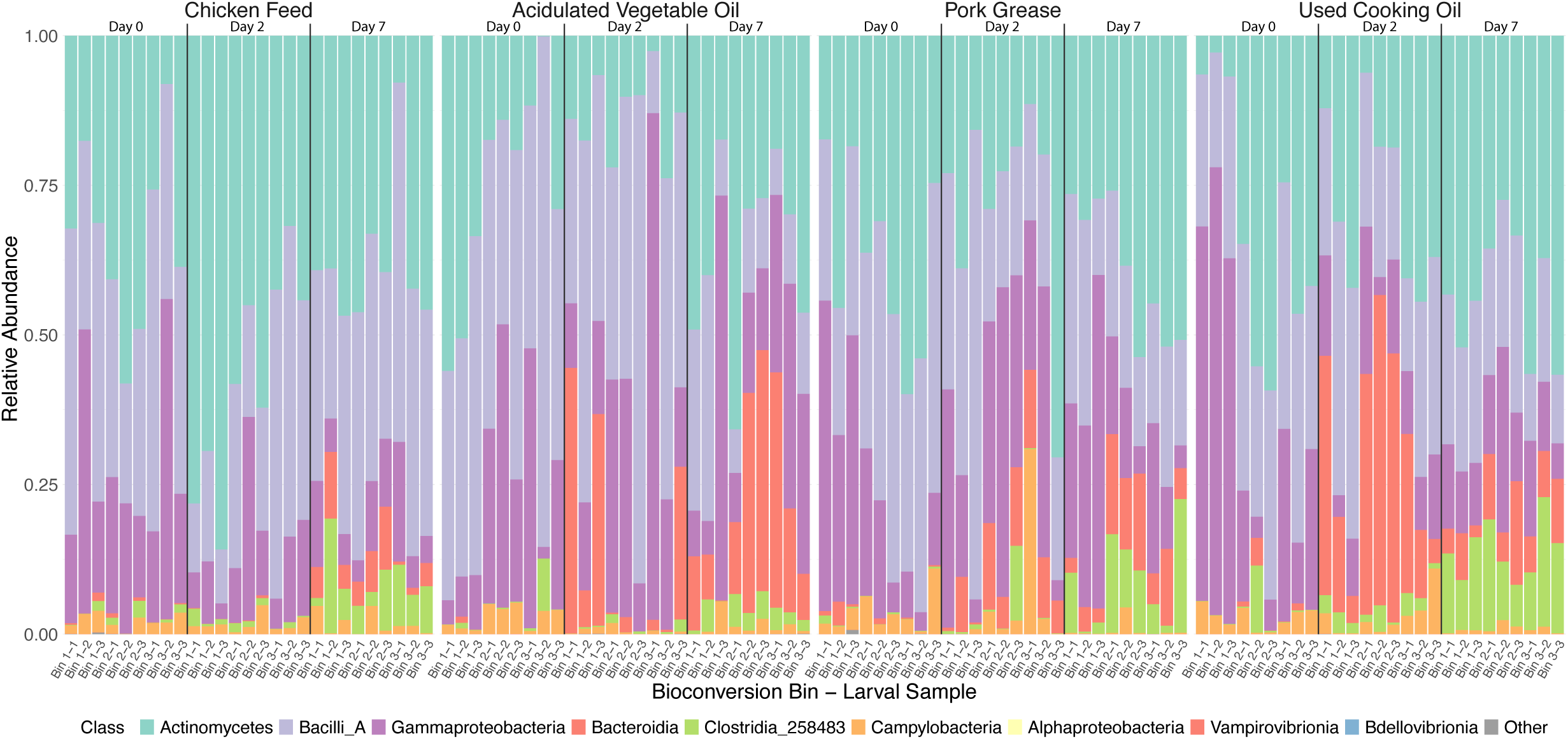
Oil supplementation impacts dominant taxa in the gut microbiome. Relative abundance plots showing class-level relative abundances of each sample by diet across bioconversion days. Colors indicate bacterial classes. All classes outside of the top 9 classes were summed as “Other”. CF = chicken feed control diet, PG = pork grease, UCO = used cooking oil, AVO = acidulated vegetable oil.

### Waste Oil Streams Differentially Impact Core BSFL Gut Microbiota

Core feature analysis identified 60 ASVs as members of the baseline core BSFL gut microbiota at bioconversion day 0 (**Table S7**). To examine how waste oil supplementation affected these and other gut community members, differential abundance analysis was conducted at the ASV level by comparing each oil-supplemented diet to the CF control at each bioconversion day. Across all comparisons, 29 unique ASVs were differentially abundant in at least one oil-supplemented treatment or timepoint, including 16 ASVs in AVO-fed larvae, 14 in PG-fed larvae, and 18 in UCO-fed larvae (**Figure 4**). Most of these differentially abundant ASVs were members of the baseline core community, with 20 of the 29 ASVs (∼69%) identified as core features at day 0 (**Table S7**).

**Figure 4.**
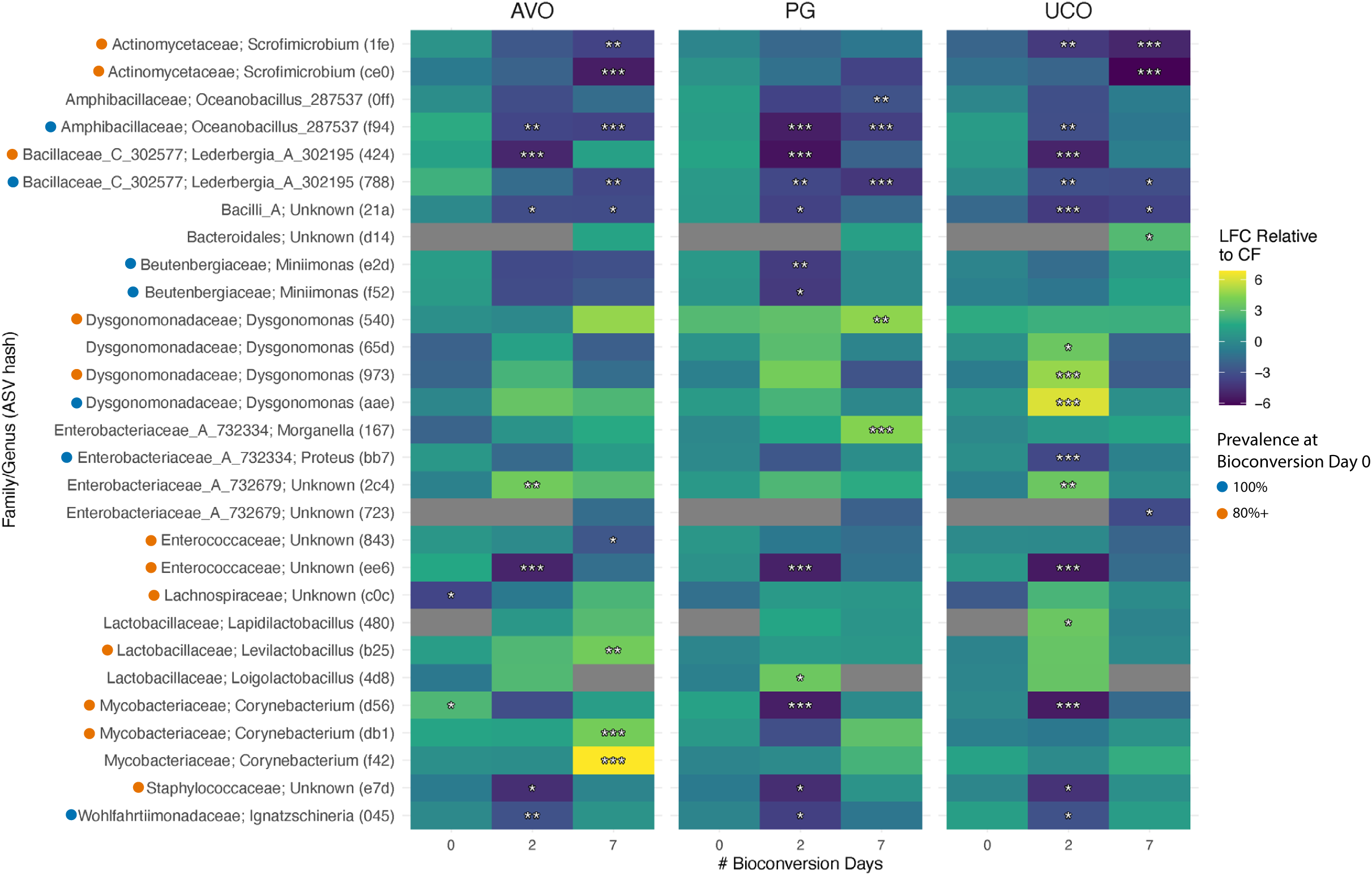
Oil supplementation enriches specific core BSFL gut microbiota. Heatmaps show differentially abundant taxa in each oil-supplemented diet relative to CF. Only ASVs with at least one significant pairwise difference were included, and are classified at the family & genus level unless the ASV was unclassified at those levels. The first three letters of the ASV hash are included in parentheses to provide a unique identifier as multiple ASVs map to the same genus. Significant differences are denoted by asterisks (*p < 0.05, **p < 0.01, ***p < 0.001). Colors indicate log fold change relative to CF. Gray boxes indicate that an ASV was not detected at all on that bioconversion day. Colored circles next to taxa indicate level of prevalence among the day 0 larvae.

Consistent with the diversity and class-level relative abundance patterns (**Figures 2**, **3**), ASV-level differences were most pronounced on day 2 (**Figure 4**). By day 7, fewer ASVs remained differentially abundant relative to CF, indicating that several oil-associated taxonomic shifts were strongest during the intermediate stage of bioconversion. ASVs assigned to *Scrofimicrobium*, *Oceanobacillus*, *Lederbergia*, *Proteus*, Enterococcaceae, *Miniimonas*, Staphylococcaceae, and *Ignatzschineria* were relatively lower in oil-supplemented diets relative to CF. ASVs assigned to *Dysgonomonas*, Lactobacillaceae, and *Morganella* were relatively higher in oil-supplemented diets. Several genera contained multiple ASVs with different response patterns across diets and timepoints. For example, ASVs assigned to *Corynebacterium* and *Dysgonomonas* did not respond uniformly, with some ASVs increasing in oil-supplemented diets relative to CF and others decreasing or remaining similar to the control.

### Most Waste Oils Maintain Larval Growth and Bioconversion Performance

No significant differences in larval growth or bioconversion traits were observed at the end of the nursery phase; however, variability was observed between bins (**Figure S4**, **Table S8-9**). Following the 7-day bioconversion, percent larval mass gain and percent diet mass loss were generally negatively impacted in the AVO-fed larvae (**Figure 5A-B**). This is highlighted by the significant reduction in bioconversion rate during the AVO treatment (**Figure 5C**). PG- and UCO-supplemented diets did not differ significantly from the CF control (**Figure 5, Table S9**).

**Figure 5.**
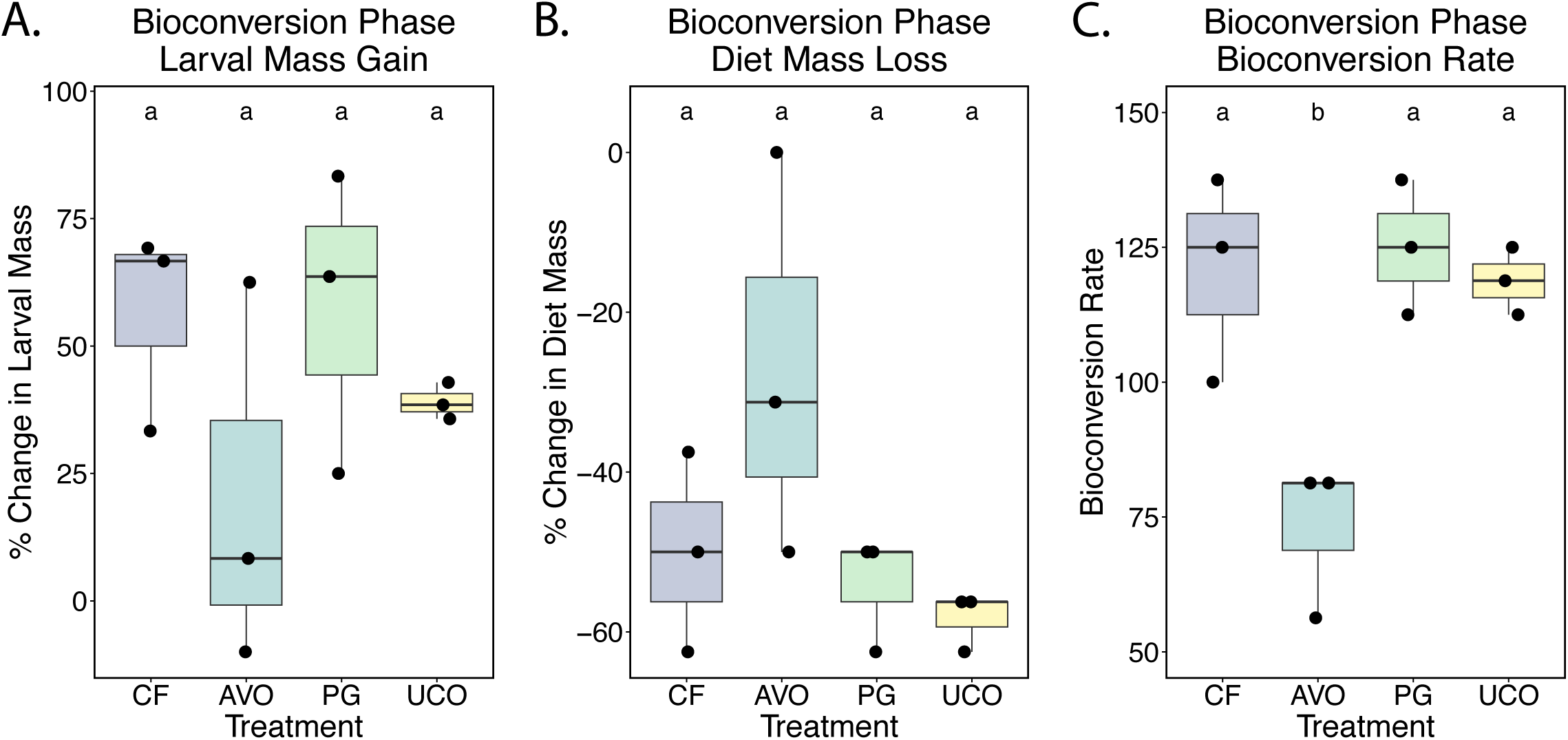
Growth and bioconversion traits of BSFL after bioconversion. Boxplots demonstrate change in (A) percent larval mass gain, (B) percent diet mass loss, and (C) bioconversion rate over the course of 7 days. Colors denote diet. CF = chicken feed control diet, PG = pork grease, UCO = used cooking oil, AVO = acidulated vegetable oil. Data were analyzed using one-way ANOVA with a post-hoc Tukey’s HSD pairwise comparisons test. Letters represent significantly different groups (p < 0.05).

### Waste Oil Substrates Alter Larval Fat and Protein Content and Modify Fatty Acid Profiles

The nutritional composition of the larvae after 7 days of bioconversion varied by diet. Total fat and protein content differed among diet groups (**Figure 6A-B**, **Table S10**). Larvae reared on oil-supplemented diets generally had higher fat content and lower protein content than CF-fed larvae, but significant pairwise differences were observed only between the CF- and UCO-fed larvae. Fat content was highest in UCO-fed larvae (28.6 ± 6.05%) and lowest in CF-fed larvae (8.01 ± 3.00%). Conversely, protein content was highest in CF-fed larvae (50.1 ± 0.962%) and lowest in UCO-fed larvae (32.2 ± 2.12%).

**Figure 6.**
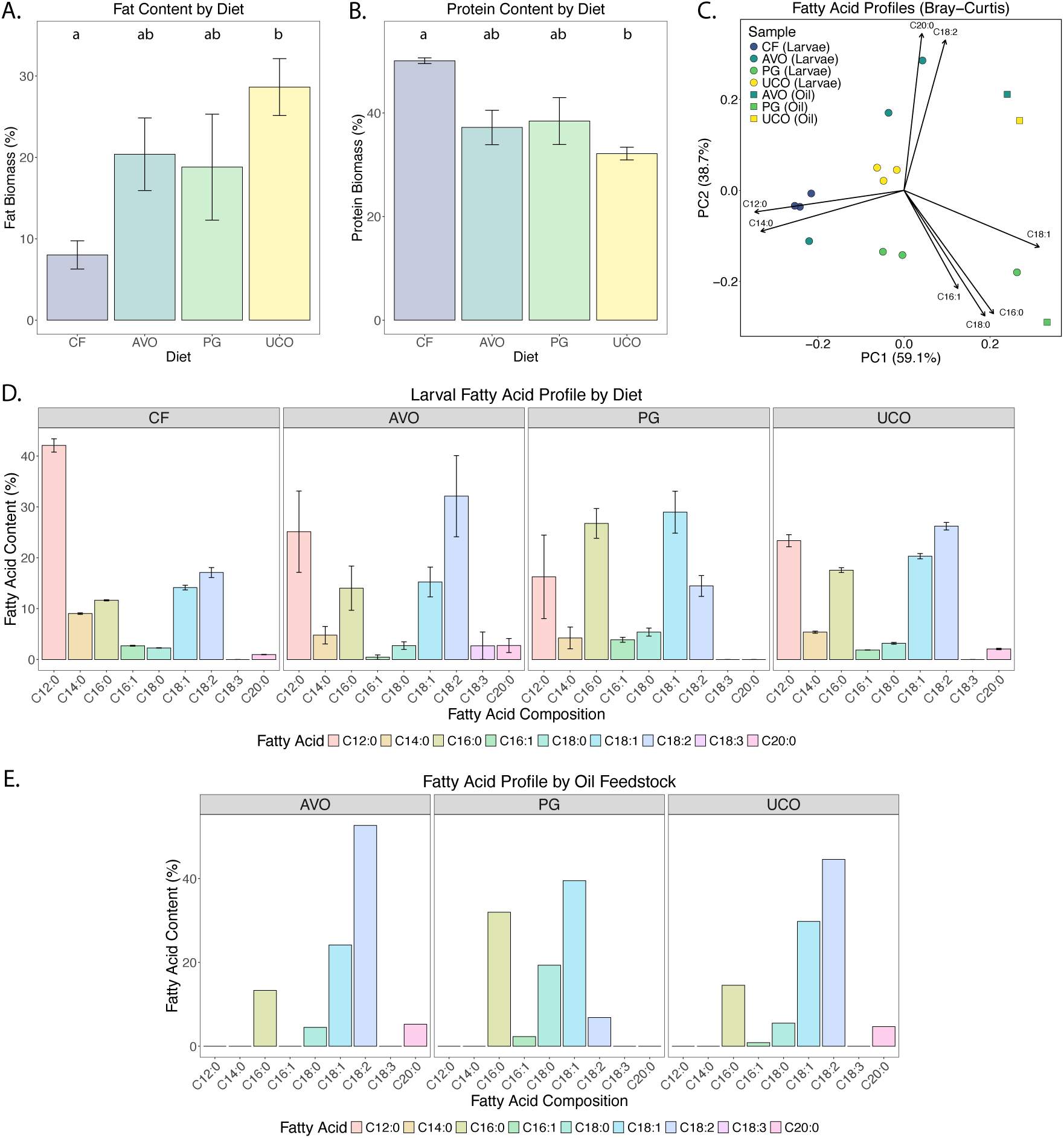
Oil supplementation alters larval biomass composition and fatty acid profiles. Larval biomass composition measured as (A) total fat content and (B) total protein content after 7 days of bioconversion. Letters denote significantly different groups (one-way ANOVA with Tukey’s HSD; p < 0.05). (C) Biplot ordination of Bray-Curtis dissimilarities calculated from larval fatty acid profiles. Points are colored by diet; square symbols indicate supplemented oil feedstock profiles included for visual comparison. Arrows indicate significant vector correlations with principal components calculated on fatty acid relative abundances. Vector lengths are scaled by their correlation strength so that “weak” predictors have shorter arrows than “strong” predictors. PERMANOVA was performed on larval samples only. Fatty acid profiles of (D) larvae and (E) supplemented oil feedstocks. Colors denote individual fatty acids, and error bars represent standard error. CF = chicken feed control, AVO = acidulated vegetable oil, PG = pork grease, and UCO = used cooking oil.

Larval fatty acid composition also differed significantly among diets (Bray-Curtis R^2^ = 0.58; p = 0.006; **Figure 6C, Table S11**). AVO-, PG-, and UCO-fed larvae each shifted toward the fatty acid composition of their corresponding oil feedstocks but occupied intermediate positions between the feedstock profiles and the CF larval profile. These spatial patterns corresponded to shifts in the relative abundances of specific fatty acids in both the larvae and supplemented oil feedstocks (**Figure 6D-E**). CF-fed larvae had the highest relative abundances of the medium-chain saturated fatty acids lauric acid (C12:0; 42.10 ± 2.26%) and myristic acid (C14:0; 9.04 ± 0.26%). Although lauric acid and myristic acid were not detected in the supplemented oil feedstocks, both fatty acids were retained in oil-supplemented larvae at lower relative abundances than in CF-fed larvae. Oil-supplemented larvae also had higher relative abundances of longer-chain fatty acids that were abundant in their corresponding feedstocks, with the largest diet-associated shifts observed in polyunsaturated fatty acids. AVO- and UCO-fed larvae had higher relative abundances of linoleic acid (C18:2), which was abundant in both AVO and UCO feedstocks. PG-fed larvae had higher relative abundances of palmitic acid (C16:0) and oleic acid (C18:1), the dominant fatty acids in pork grease; however, the elevated stearic acid (C18:0) present in the feedstock was not proportionally reflected in the larvae.

## DISCUSSION

This study demonstrates that lipid-rich waste oil streams can reshape BSFL gut microbiome membership, abundance-weighted community structure, and larval nutritional composition in a substrate-specific manner. Larval age was a major driver of gut microbiome differentiation, consistent with previous studies showing that the BSFL gut microbiome changes across larval development and life stages (14–17, 42, 43). Waste oil supplementation also altered gut community composition, with diet-associated effects becoming strongest during early-to-intermediate bioconversion. However, microbial restructuring did not translate uniformly to larval performance. PG- and UCO-supplemented diets maintained larval growth and bioconversion performance comparable to the CF control, whereas AVO significantly reduced bioconversion rate and showed a non-significant trend toward lower larval mass. These results show that some waste oil streams can be incorporated into BSFL systems without compromising larval performance while modifying gut microbiome structure and larval biomass composition.

Although oil supplementation altered gut community composition, alpha diversity did not show a consistent response. Shannon diversity, evenness, and phylogenetic diversity varied across timepoints and diets, indicating that lipid-rich waste oils did not uniformly increase or decrease within-sample diversity. Instead, the strongest effects involved changes in community membership and relative abundance, suggesting substrate-specific filtering of gut community members. Class-level patterns supported this interpretation, since during early-to-intermediate bioconversion, oil-supplemented diets diverged from the Actinomycetes-dominated trajectory observed in CF, whereas later increases in Clostridia across treatments may reflect shared developmental or late-stage bioconversion processes. Day 0 differences were primarily membership-based and accompanied by strong bin effects, suggesting that initial variation among larvae or rearing bins may have reflected low-abundance taxa, priority effects, or stochastic colonization before diet-driven filtering became stronger. This is consistent with gut and decomposer systems in which early stochastic assembly can influence initial community structure, while deterministic selection increases as substrate chemistry, nutrient availability, and microbial interactions change during decomposition (44, 45).

At finer taxonomic resolution, ASV-level patterns showed that much of the substrate-associated response involved members of the baseline core gut microbiome. Most differentially abundant ASVs were present before bioconversion, suggesting that oil supplementation altered the relative success of resident taxa while also contributing to substrate-specific turnover. This resident community reweighting is consistent with the role of BSFL as generalist decomposers (1, 2, 46), whose broad dietary range may be supported by gut microbiome plasticity (15, 47, 48). Our results extend this to lipid-rich waste oil streams, showing that the BSFL gut microbiome can respond to oil substrates that are difficult to incorporate into conventional waste valorization pipelines (23). However, ASV responses were not always consistent within genera. Several genera contained ASVs with contrasting responses across diets and timepoints, suggesting that genus-level interpretation may obscure ecologically meaningful variation among closely related community members. Consequently, enrichment or depletion of oil-responsive taxa should be interpreted as evidence of ecological responsiveness rather than a single shared metabolic role. For example, enrichment of *Dysgonomonas*, *Corynebacterium*, and *Morganella* may reflect lipid tolerance, passive responses to altered gut conditions, indirect associations with other degraders, or direct involvement in lipid transformation and nutrient assimilation. Prior associations of *Dysgonomonas* and *Corynebacterium* with complex substrate metabolism, such as lipids (14, 49), plastics (50–53), and lignin (54, 55), and of *Morganella* with vertebrate decomposition environments (44), support their potential relevance to organic waste bioconversion. In contrast, *Ignatzschneria* was depleted across all three oil-supplemented diets, despite being commonly associated with flies and decomposition environments (56). Because *Ignatzschneria* has often been linked to decomposition of more labile organic resources (44), its consistent depletion may indicate that waste oil substrates shift gut conditions away from conventional decomposition-associated communities and toward communities better suited to oil waste.

Despite microbial responses across all oil treatments, larval performance varied among feedstocks, indicating that waste oil effects were not driven by total lipid content alone. The three oil substrates differed in characteristics likely to influence host physiology and microbial activity: PG was more saturated, AVO was more acidic and enriched in free fatty acids (FFAs), and repeatedly heated UCOs typically contain elevated degradation products from lipid hydrolysis, oxidation, and polymerization, including FFAs, oxidized compounds, and polymerized acylglycerides (23). Although these substrates produced distinct microbiome responses, the specific chemical features driving those shifts remain unknown. PG- and UCO-supplemented diets supported larval growth and bioconversion performance comparable to CF, whereas AVO reduced bioconversion rate and showed weaker growth outcomes. This contrast indicates that microbiome restructuring alone does not predict successful bioconversion; substrate value depends on whether larvae can maintain feeding, growth, and lipid assimilation under a given chemical environment. AVO likely presented a distinct challenge because acidulated oils are enriched in FFAs (22), increasing acidity and potentially disrupting membranes, lipid oxidation dynamics, and microbial growth (57, 58). In contrast, PG and UCO may have remained more compatible with larval feeding, digestion, and lipid assimilation despite differences in saturation and thermal degradation history. Observations of AVO-fed larvae climbing container walls rather than burrowing into the substrate were consistent with substrate avoidance or stress, although weaker nursery-phase growth trends mean pre-existing host conditions cannot be excluded. Reduced AVO performance may therefore reflect a combination of substrate-specific stress, altered feeding behavior, initial larval condition, and/or inhibition of microbial taxa involved in nutrient transformation. Further work is needed to determine whether AVO can be incorporated more effectively through lower inclusion rates, pH neutralization, or other pretreatment approaches.

Prior work has shown that rearing substrate can influence BSFL biomass composition (59), although protein and fat responses may depend on broader macronutrient balance, developmental stage, and substrate quality rather than substrate composition alone. Our nutritional analyses suggest that oil supplementation altered larval biomass through dietary lipid incorporation, host metabolism, and potentially microbial processing. Oil-fed larvae generally had higher fat content and lower protein content than CF-fed larvae, with the strongest pairwise differences observed between CF- and UCO-fed larvae. This suggests that BSFL can accumulate substrate-derived lipids, whereas protein content may be more constrained by larval physiology or broader substrate macronutrient balance, including protein-to-carbohydrate ratios (60). Fatty-acid profiles provided clearer evidence that oil substrates shaped larval biomass composition. Oil-fed larvae shifted toward the fatty-acid composition of their corresponding feedstocks, consistent with the broader pattern that insect biomass composition can reflect dietary fatty-acid inputs, including in BSFL (59). However, larval profiles remained intermediate between the oil feedstocks and CF-fed larvae, indicating that biomass composition was shaped by both dietary input and biological processing. The persistence of lauric and myristic acids in oil-fed larvae, despite their absence from the supplemented oil fractions, likely reflects contributions from the basal chicken feed, larval lipid metabolism, and/or microbial transformations. Likewise, enrichment of linoleic acid in AVO- and UCO-fed larvae and palmitic/oleic acids in PG-fed larvae indicates incorporation of feedstock-enriched fatty acids, whereas incomplete transfer of stearic acid in PG-fed larvae may reflect selective assimilation, metabolism, oxidation, or differential retention. Because linoleic acid is an essential omega-6 polyunsaturated fatty acid for humans and many livestock animals, its enrichment may be relevant for alternative food and feed applications in agriculture (61, 62). These results suggest that lipid-rich waste products can serve as both energy-dense substrates and tools for shaping larval fatty-acid profiles, converting low-value lipid by-products into higher-value biomass while retaining the influence of basal diet and larval metabolism.

Overall, this study highlights BSFL bioconversion as a host-microbiome agricultural biotechnology in which lipid-rich agricultural and food-processing by-products can be transformed into insect biomass with substrate-dependent nutritional profiles. Waste oil substrates redirected gut microbiome assembly and altered larval fatty-acid composition, but substrate identity determined whether these changes were compatible with efficient bioconversion. PG and UCO supported larval performance while modifying biomass composition, whereas AVO showed that substrate chemistry can constrain conversion efficiency and may require optimization before use. These findings indicate that BSFL production outcomes are shaped by interactions among gut community plasticity, larval physiology, and waste-stream chemistry, which together influence both bioconversion performance and final biomass composition. Because this study used 16S rRNA gene sequencing and endpoint nutritional analyses, future work should use chemically defined substrate experiments to isolate how lipid saturation, free fatty acids, acidity, oxidation products, polar compounds, and polymerized acylglycerols influence larval growth, gut microbiome function, and lipid assimilation. Pairing these trials with metagenomic, metatranscriptomic, metabolomic, and substrate-optimization approaches would help identify microbial pathways and substrate features that promote efficient lipid-waste conversion. By showing that PG and UCO can support larval performance while restructuring the gut microbiome and host fatty-acid composition, this study demonstrates how BSFL-microbiome systems could contribute to a circular agricultural bioeconomy by converting underutilized lipid wastes into value-added insect biomass.

## ACKNOWLEDGMENTS

This work was partially funded from start-up funds provided to Z.B. from the University of Tennessee, the Oak Ridge Associated Universities-Directed Research Development grant and the University of Tennessee AgResearch Seed Grant awarded to C.G.O, and the National Institute of Food and Agriculture as part of the United States Department of Agriculture (Grant No. 13987835) award to T.W. R.S. was supported by the University of Tennessee-Oak Ridge Innovation Institute Graduate Student Fellowship during the project period. This research received no specific grant from any funding agency in the public, commercial, or not-for-profit sectors. The funders had no role in study design, data collection and interpretation, or the decision to submit the work for publication.

## REFERENCES

1. Tomberlin JK, Sheppard DC, Joyce JA. 2002. Selected Life-History Traits of Black Soldier Flies (Diptera: Stratiomyidae) Reared on Three Artificial Diets. Ann Entomol Soc Am 95:379–386.

2. Liu T, Klammsteiner T, Dregulo AM, Kumar V, Zhou Y, Zhang Z, Awasthi MK. 2022. Black soldier fly larvae for organic manure recycling and its potential for a circular bioeconomy: A review. Sci Total Environ 833:155122.

3. Bruno D, Orlando M, Testa E, Carnevale Miino M, Pesaro G, Miceli M, Pollegioni L, Barbera V, Fasoli E, Draghi L, Baltrocchi APD, Ferronato N, Seri R, Maggi E, Caccia S, Casartelli M, Molla G, Galimberti MS, Torretta V, Vezzulli A, Tettamanti G. 2025. Valorization of organic waste through black soldier fly: On the way of a real circular bioeconomy process. Waste Manag 191:123–134.

4. Su H, Zhang B, Shi J, He S, Dai S, Zhao Z, Wu D, Li J. 2025. Black Soldier Fly Larvae as a Novel Protein Feed Resource Promoting Circular Economy in Agriculture. Insects 16:830.

5. Liu R, Qu X, Gao Z, Luan Y, Zheng Y, Wu L, Shang T, Teng T, Shi B. 2026. Bacillus velezensis enhances the conversion of swine manure into high-quality insect protein via augmented histidine metabolism in Hermetia illucens larvae. Bioresour Technol 455:134865.

6. Bruno D, Casartelli M, De Smet J, Gold M, Tettamanti G. 2025. Review: A journey into the black soldier fly digestive system: From current knowledge to applied perspectives. animal 19:101483.

7. Yu Z, Xie C, Zhang Z, Huang Z, Zhou J, Wang C. 2024. Microbial fermentation and black soldier fly feeding to enhance maize straw degradation. Chemosphere 353:141498.

8. Xiang F, Zhang Q, Xu X, Zhang Z. 2024. Black soldier fly larvae recruit functional microbiota into the intestines and residues to promote lignocellulosic degradation in domestic biodegradable waste. Environ Pollut 340:122676.

9. De Filippis F, Bonelli M, Bruno D, Sequino G, Montali A, Reguzzoni M, Pasolli E, Savy D, Cangemi S, Cozzolino V, Tettamanti G, Ercolini D, Casartelli M, Caccia S. 2023. Plastics shape the black soldier fly larvae gut microbiome and select for biodegrading functions. Microbiome 11:205.

10. Dragone NB, van Hamelsveld S, Nazmi AR, Stott M, Hatley GA, Moloney K, Bohm K, Gutierrez-Gines MJ, Weaver L. 2025. Examining the potential of plastic-fed black soldier fly larvae (Hermetia illucens) as “bioincubators” of plastic-degrading bacteria. J Appl Microbiol 136:lxaf085.

11. Bruno D, Bonelli M, De Filippis F, Di Lelio I, Tettamanti G, Casartelli M, Ercolini D, Caccia S. 2019. The Intestinal Microbiota of Hermetia illucens Larvae Is Affected by Diet and Shows a Diverse Composition in the Different Midgut Regions. Appl Environ Microbiol 85:e01864–18.

12. Pei Y, Zhao S, Chen X, Zhang J, Ni H, Sun M, Lin H, Liu X, Chen H, Yang S. 2022. Bacillus velezensis EEAM 10B Strengthens Nutrient Metabolic Process in Black Soldier Fly Larvae (Hermetia illucens) via Changing Gut Microbiome and Metabolic Pathways. Front Nutr 9:880488.

13. Auger L, Deschamps M-H, Vandenberg G, Derome N. 2025. First Attempt at Synthetic Microbial Communities Design for Rearing Gnotobiotic Black Soldier Fly Hermetia illucens (Linnaeus) Larvae. Insects 16:851.

14. Klammsteiner T, Walter A, Bogataj T, Heussler CD, Stres B, Steiner FM, Schlick-Steiner BC, Insam H. 2021. Impact of Processed Food (Canteen and Oil Wastes) on the Development of Black Soldier Fly (Hermetia illucens) Larvae and Their Gut Microbiome Functions. Front Microbiol 12:619112.

15. Bruno D, Bonelli M, De Filippis F, Di Lelio I, Tettamanti G, Casartelli M, Ercolini D, Caccia S. 2019. The Intestinal Microbiota of Hermetia illucens Larvae Is Affected by Diet and Shows a Diverse Composition in the Different Midgut Regions. Appl Environ Microbiol 85:e01864–18.

16. Shelomi M, Wu M-K, Chen S-M, Huang J-J, Burke CG. 2020. Microbes Associated With Black Soldier Fly (Diptera: Stratiomiidae) Degradation of Food Waste. Environ Entomol 49:405–411.

17. Xu L, Lin Q, Wang S, Chen S, Yang R, Liu C, Hu Q, Zhao Z, Cao Z. 2025. Efficacy of black soldier fly larvae in converting kitchen waste and the dynamic alterations of their gut microbiome. J Environ Manage 377:124613.

18. IJdema F, De Smet J, Crauwels S, Lievens B, Van Campenhout L. 2022. Meta-analysis of larvae of the black soldier fly (Hermetia illucens) microbiota based on 16S rRNA gene amplicon sequencing. FEMS Microbiol Ecol 98:fiac094.

19. Klammsteiner T, Walter A, Bogataj T, Heussler CD, Stres B, Steiner FM, Schlick-Steiner BC, Arthofer W, Insam H. 2020. The Core Gut Microbiome of Black Soldier Fly (Hermetia illucens) Larvae Raised on Low-Bioburden Diets. Front Microbiol 11:993.

20. Koch H, Lessard B, Escobar-Correas S, Thurman JH, Paten AM, Morgan MJ. 2026. A Comparative Survey of Soldier Fly (Stratiomyidae) Larval Gut Microbiomes Across Five Subfamilies Reveals Novel Bacterial Diversity and a “Wild Core” in Hermetia illucens. Microb Ecol 10.1007/s00248-026-02760-z.

21. Gold M, von Allmen F, Zurbrügg C, Zhang J, Mathys A. 2020. Identification of Bacteria in Two Food Waste Black Soldier Fly Larvae Rearing Residues. Front Microbiol 11.

22. Casali B, Brenna E, Parmeggiani F, Tessaro D, Tentori F. 2021. Enzymatic Methods for the Manipulation and Valorization of Soapstock from Vegetable Oil Refining Processes. Sustain Chem 2:74–91.

23. Beghetto V. 2025. Strategies for the Transformation of Waste Cooking Oils into High-Value Products: A Critical Review. Polymers 17:368.

24. Choe E, Min D b. 2007. Chemistry of Deep-Fat Frying Oils. J Food Sci 72:R77–R86.

25. Kerr BJ, Lindblom SC, Zhao J, Faris RJ. 2020. Influence of feeding thermally peroxidized lipids on growth performance, lipid digestibility, and oxidative status in nursery pigs. J Anim Sci 98:skaa392.

26. Kumar A, Bhayana S, Singh PK, Tripathi AD, Paul V, Balodi V, Agarwal A. 2025. Valorization of used cooking oil: challenges, current developments, life cycle assessment and future prospects. Discov Sustain 6:119.

27. Lotero E, Liu Y, Lopez DE, Suwannakarn K, Bruce DA, Goodwin JG. 2005. Synthesis of Biodiesel via Acid Catalysis. Ind Eng Chem Res 44:5353–5363.

28. Bolyen E, Rideout JR, Dillon MR, Bokulich NA, Abnet CC, Al-Ghalith GA, Alexander H, Alm EJ, Arumugam M, Asnicar F, Bai Y, Bisanz JE, Bittinger K, Brejnrod A, Brislawn CJ, Brown CT, Callahan BJ, Caraballo-Rodríguez AM, Chase J, Cope EK, Da Silva R, Diener C, Dorrestein PC, Douglas GM, Durall DM, Duvallet C, Edwardson CF, Ernst M, Estaki M, Fouquier J, Gauglitz JM, Gibbons SM, Gibson DL, Gonzalez A, Gorlick K, Guo J, Hillmann B, Holmes S, Holste H, Huttenhower C, Huttley GA, Janssen S, Jarmusch AK, Jiang L, Kaehler BD, Kang KB, Keefe CR, Keim P, Kelley ST, Knights D, Koester I, Kosciolek T, Kreps J, Langille MGI, Lee J, Ley R, Liu Y-X, Loftfield E, Lozupone C, Maher M, Marotz C, Martin BD, McDonald D, McIver LJ, Melnik AV, Metcalf JL, Morgan SC, Morton JT, Naimey AT, Navas-Molina JA, Nothias LF, Orchanian SB, Pearson T, Peoples SL, Petras D, Preuss ML, Pruesse E, Rasmussen LB, Rivers A, Robeson MS, Rosenthal P, Segata N, Shaffer M, Shiffer A, Sinha R, Song SJ, Spear JR, Swafford AD, Thompson LR, Torres PJ, Trinh P, Tripathi A, Turnbaugh PJ, Ul-Hasan S, van der Hooft JJJ, Vargas F, Vázquez-Baeza Y, Vogtmann E, von Hippel M, Walters W, Wan Y, Wang M, Warren J, Weber KC, Williamson CHD, Willis AD, Xu ZZ, Zaneveld JR, Zhang Y, Zhu Q, Knight R, Caporaso JG. 2019. Reproducible, interactive, scalable and extensible microbiome data science using QIIME 2. Nat Biotechnol 37:852–857.

29. Callahan BJ, McMurdie PJ, Rosen MJ, Han AW, Johnson AJA, Holmes SP. 2016. DADA2: High resolution sample inference from Illumina amplicon data. Nat Methods 13:581–583.

30. Davis NM, Proctor DM, Holmes SP, Relman DA, Callahan BJ. 2018. Simple statistical identification and removal of contaminant sequences in marker-gene and metagenomics data. Microbiome 6:226.

31. Glöckner FO, Yilmaz P, Quast C, Gerken J, Beccati A, Ciuprina A, Bruns G, Yarza P, Peplies J, Westram R, Ludwig W. 2017. 25 years of serving the community with ribosomal RNA gene reference databases and tools. J Biotechnol 261:169–176.

32. Bokulich NA, Kaehler BD, Rideout JR, Dillon M, Bolyen E, Knight R, Huttley GA, Gregory Caporaso J. 2018. Optimizing taxonomic classification of marker-gene amplicon sequences with QIIME 2’s q2-feature-classifier plugin. Microbiome 6:90.

33. McDonald D, Jiang Y, Balaban M, Cantrell K, Zhu Q, Gonzalez A, Morton JT, Nicolaou G, Parks DH, Karst SM, Albertsen M, Hugenholtz P, DeSantis T, Song SJ, Bartko A, Havulinna AS, Jousilahti P, Cheng S, Inouye M, Niiranen T, Jain M, Salomaa V, Lahti L, Mirarab S, Knight R. 2024. Greengenes2 unifies microbial data in a single reference tree. Nat Biotechnol 42:715–718.

34. Anderson MJ. 2017. Permutational Multivariate Analysis of Variance (PERMANOVA), p. 1–15. In Wiley StatsRef: Statistics Reference Online. John Wiley & Sons, Ltd.

35. R Core Team. 2026. R: A Language and Environment for Statistical Computing (4.4.2). R Foundation for Statistical Computing, Vienna, Austria.

36. Folch J, Lees M, Stanley GHS. 1957. A SIMPLE METHOD FOR THE ISOLATION AND PURIFICATION OF TOTAL LIPIDES FROM ANIMAL TISSUES. J Biol Chem 226:497–509.

37. Ismail BP, Nielsen SS. 2024. Nielsen’s Food Analysis. Springer International Publishing, Cham. https://link.springer.com/10.1007/978-3-031-50643-7. Retrieved 28 April 2026.

38. Yuan Y, Trinh D, Hoesel S, Jagadamma S, DeBruyn J, Abdoumoumine N, Wang T. 2026. Black soldier fly larvae drying methods affect Osborne protein fractionation. Appl Food Res 6:101767.

39. Eder K. 1995. Gas chromatographic analysis of fatty acid methyl esters. J Chromatogr B Biomed Appl 671:113–131.

40. Angeletti B, Trinh DT, Dia V, Burns S, Chester MA, Bergee RE, Wang T. 2025. Hempseed Hydrolysates Exhibit Antioxidant Activity in Meat Systems. Foods 14:1728.

41. Oksanen J, Simpson G, Blanchet F, Kindt R, Legendre P, Minchin P, O’Hara R, Solymos P, Stevens M, Szoecs E, Wagner H, Barbour M, Bedward M, Bolker B, Borcard D, Borman T, Carvalho G, Chirico M, De Caceres M, Durand S, Evangelista H, FitzJohn R, Friendly M, Furneaux B, Hannigan G, Hill M, Lahti L,, Martino C, McGlinn D, Ouellette M, Ribeiro Cunha E, Smith T, Stier A, Ter Braak C, Weedon J. 2025. vegan: Community Ecology Package (2.7.1).

42. Li X-Y, Mei C, Luo X-Y, Wulamu D, Zhan S, Huang Y-P, Yang H. 2023. Dynamics of the intestinal bacterial community in black soldier fly larval guts and its influence on insect growth and development. Insect Sci 30:947–963.

43. Klammsteiner T, Heussler CD, Stonig KT, Insam H, Schlick-Steiner BC, Steiner FM. 2026. Stage-Specific Microbiota Transitions Throughout Black Soldier Fly Ontogeny. Microb Ecol 89:41.

44. Burcham ZM, Belk AD, McGivern BB, Bouslimani A, Ghadermazi P, Martino C, Shenhav L, Zhang AR, Shi P, Emmons A, Deel HL, Xu ZZ, Nieciecki V, Zhu Q, Shaffer M, Panitchpakdi M, Weldon KC, Cantrell K, Ben-Hur A, Reed SC, Humphry GC, Ackermann G, McDonald D, Chan SHJ, Connor M, Boyd D, Smith J, Watson JMS, Vidoli G, Steadman D, Lynne AM, Bucheli S, Dorrestein PC, Wrighton KC, Carter DO, Knight R, Metcalf JL. 2024. A conserved interdomain microbial network underpins cadaver decomposition despite environmental variables. Nat Microbiol 9:595–613.

45. Jones EW, Carlson JM, Sivak DA, Ludington WB. 2022. Stochastic microbiome assembly depends on context. Proc Natl Acad Sci 119:e2115877119.

46. Diener S, Zurbrügg C, Tockner K. 2009. Conversion of organic material by black soldier fly larvae: establishing optimal feeding rates. Waste Manag Res 27:603–610.

47. Tanga CM, Waweru JW, Tola YH, Onyoni AA, Khamis FM, Ekesi S, Paredes JC. 2021. Organic Waste Substrates Induce Important Shifts in Gut Microbiota of Black Soldier Fly (Hermetia illucens L.): Coexistence of Conserved, Variable, and Potential Pathogenic Microbes. Front Microbiol 12:635881.

48. Dragone NB, van Hamelsveld S, Nazmi AR, Stott M, Hatley GA, Moloney K, Bohm K, Gutierrez-Gines MJ, Weaver L. 2025. Examining the potential of plastic-fed black soldier fly larvae (Hermetia illucens) as “bioincubators” of plastic-degrading bacteria. J Appl Microbiol 136:lxaf085.

49. Tauch A, Kaiser O, Hain T, Goesmann A, Weisshaar B, Albersmeier A, Bekel T, Bischoff N, Brune I, Chakraborty T, Kalinowski J, Meyer F, Rupp O, Schneiker S, Viehoever P, Pühler A. 2005. Complete genome sequence and analysis of the multiresistant nosocomial pathogen Corynebacterium jeikeium K411, a lipid-requiring bacterium of the human skin flora. J Bacteriol 187:4671–4682.

50. Wang Y, Luo L, Li X, Wang J, Wang H, Chen C, Guo H, Han T, Zhou A, Zhao X. 2022. Different plastics ingestion preferences and efficiencies of superworm (*Zophobas atratus* Fab.) and yellow mealworm (*Tenebrio molitor* Linn.) associated with distinct gut microbiome changes. Sci Total Environ 837:155719.

51. Sun J, Prabhu A, Aroney STN, Rinke C. 2022. Insights into plastic biodegradation: community composition and functional capabilities of the superworm (Zophobas morio) microbiome in styrofoam feeding trials. Microb Genomics 8:000842.

52. Su Y, Zhang Z, Zhu J, Shi J, Wei H, Xie B, Shi H. 2021. Microplastics act as vectors for antibiotic resistance genes in landfill leachate: The enhanced roles of the long-term aging process. Environ Pollut 270:116278.

53. Luo L, Wang Y, Guo H, Yang Y, Qi N, Zhao X, Gao S, Zhou A. 2021. Biodegradation of foam plastics by *Zophobas atratus* larvae (Coleoptera: Tenebrionidae) associated with changes of gut digestive enzymes activities and microbiome. Chemosphere 282:131006.

54. Bridges CM, Gage DJ. 2021. Draft Genome Sequences of Dysgonomonas sp. Strains GY75 and GY617, Isolated from the Hindgut of Reticulitermes flavipes. Microbiol Resour Announc 10:e00079–21.

55. Duan J, Liang J, Wang Y, Du W, Wang D. 2016. Kraft Lignin Biodegradation by Dysgonomonas sp. WJDL-Y1, a New Anaerobic Bacterial Strain Isolated from Sludge of a Pulp and Paper Mill 26:1765–1773.

56. Deel HL, Montoya S, King K, Emmons AL, Huhn C, Lynne AM, Metcalf JL, Bucheli SR. 2022. The microbiome of fly organs and fly-human microbial transfer during decomposition. Forensic Sci Int 340:111425.

57. Nogueira MS, Scolaro B, Milne GL, Castro IA. 2019. Oxidation products from omega-3 and omega-6 fatty acids during a simulated shelf life of edible oils. LWT 101:113–122.

58. Waraho T, McClements DJ, Decker EA. 2011. Impact of free fatty acid concentration and structure on lipid oxidation in oil-in-water emulsions. Food Chem 129:854–859.

59. Riekkinen K, Väkeväinen K, Korhonen J. 2022. The Effect of Substrate on the Nutrient Content and Fatty Acid Composition of Edible Insects. Insects 13:590.

60. Bava L, Jucker C, Gislon G, Lupi D, Savoldelli S, Zucali M, Colombini S. 2019. Rearing of Hermetia Illucens on Different Organic By-Products: Influence on Growth, Waste Reduction, and Environmental Impact. Animals 9:289.

61. Malcicka M, Visser B, Ellers J. 2018. An Evolutionary Perspective on Linoleic Acid Synthesis in Animals. Evol Biol 45:15–26.

62. Raes K, De Smet S, Demeyer D. 2004. Effect of dietary fatty acids on incorporation of long chain polyunsaturated fatty acids and conjugated linoleic acid in lamb, beef and pork meat: a review. Anim Feed Sci Technol 113:199–221.

